# Germ Granules Coordinate RNA-based Epigenetic Inheritance Pathways

**DOI:** 10.1101/707596

**Authors:** Anne E. Dodson, Scott Kennedy

## Abstract

Germ granules are biomolecular condensates that promote germ cell totipotency in most, if not all, animals. In *C. elegans*, MEG-3 and MEG-4 are two intrinsically disordered proteins that are redundantly required for the phase separations that drive germ granule assembly in germline blastomeres. Here, we show that animals lacking MEG-3/4 exhibit defects in dsRNA-mediated gene silencing (RNAi) that are due, at least in part, to defects in systemic RNAi. Interestingly, these RNAi defects are transgenerationally disconnected from *meg-3/4* genotype: RNAi defects do not arise until 5-9 generations after animals become mutant for *meg-3/4*, and RNAi defects persist for 9-11 generations after *meg-3/4* genotype is restored to wild type. Similar non-Mendelian patterns of inheritance are associated with other mutations that disrupt germ granule formation, indicating that germ granule disruption is the likely cause of genotype/phenotype disconnects. Loss of germ granules is associated with the production of aberrant populations of endogenous siRNAs, which, remarkably, are propagated for ≅10 generations in wild-type descendants of animals that lacked germ granules. *sid-1*, which encodes a factor required for systemic RNAi in *C. elegans*, is inappropriately and heritably silenced by aberrantly expressed *sid-1* endogenous siRNAs, suggesting that transgenerational silencing of *sid-1* likely underlies the heritable defect in RNAi. We conclude that one function of germ granules is to organize RNA-based epigenetic inheritance pathways and that failure to assemble germ granules has consequences that persist across many generations.

## Introduction

Cells contain many non-membrane-bound organelles (termed liquid droplet organelles or biomolecular condensates) that consist of proteins and RNAs that self-assemble via liquid-liquid phase separations^1^. Examples of biomolecular condensates include nucleoli, processing (P) bodies, Cajal bodies, stress granules, neuronal granules, and germ granules^1^. Germ granules are biomolecular condensates that form in the germ cells of many metazoans to help maintain totipotency of the germline^2^. The mechanism(s) by which germ granules promote germ cell health are largely unknown^2,3^. According to current models, one major function of biomolecular condensates such as germ granules may be to bring specific proteins and nucleic acids together in space and time to help organize the complex RNA processing pathways underlying gene regulation^4,5^.

Germ granules in *C. elegans* (referred to as P granules) are present in germ cells during all stages of development^6^. During early embryonic cell divisions, P granules assemble asymmetrically in the germline-destined portion of the zygotic cytoplasm^7^. MEG-3 and MEG-4 are intrinsically disordered proteins that are expressed during early embryogenesis and that function redundantly to nucleate P granule formation in early embryos^8,9^. Another protein, DEPS-1, contributes to P granule formation during most stages of germline development^10^. For much of development, P granules localize to the outer nuclear membrane directly adjacent to nuclear pores. Multiple lines of evidence suggest that P granules help surveil and/or process mRNAs as they transit through nuclear pores and enter the cytoplasm^11,12^. For example, newly synthesized mRNAs have been observed transiting P granules^12^, and specific mRNAs concentrate in P granules during specific stages of germline development^13^. Additionally, a number of RNA quality control proteins, including small RNA pathway components, localize to P granules (see below). Finally, in the absence of P granules, somatic genes become improperly expressed in the germline^14,15^. Thus, P granules likely function to store and surveil mRNAs in germ cells.

Non-coding RNAs, such as PIWI-interacting RNAs (piRNAs) and endogenous small interfering RNAs (endo-siRNAs), contribute to RNA quality control in many eukaryotes and, in some cases, also transmit epigenetic information from parent to progeny^16^. In *C. elegans*, many instances of transgenerational epigenetic inheritance (TEI) are directed by endo-siRNAs^17^. Current models posit that, during TEI, endo-siRNAs are deposited into the embryo via the egg or sperm. Parentally deposited endo-siRNAs then act as guide molecules to identify cognate mRNAs, recruit RNA-dependent RNA Polymerases (RdRPs), and amplify endo-siRNA populations. Repetition of this process each new generation allows endo-siRNA-based gene regulatory information to pass across multiple generations. Heritably maintained endo-siRNA populations bind Argonaute proteins such as HRDE-1 to regulate gene expression in germ cells^18,19^. Many of the genomic loci targeted for heritable silencing by endo-siRNAs in *C. elegans* are pseudogenes and cryptic loci, suggesting that the endo-siRNA system may have evolved to silence unwanted germline RNAs^20^. This heritable gene regulatory pathway is likely important for germ cell function, as mutations that disrupt this process also cause a germline mortal (Mrt) phenotype in which germ cell function deteriorates over generations^18,21,22^. In summary, *C. elegans* possess an RNA-based mode of epigenetic inheritance driven by generationally repeated amplification of endo-siRNAs by RdRPs, followed by silencing of unwanted RNAs by germline-expressed Argonautes such as HRDE-1.

Because endo-siRNA-based gene regulation is both complex and heritable, it is likely that cells regulate and organize endo-siRNA biogenesis in ways that prevent runaway heritable silencing of the wrong mRNAs. P granules may provide this organization, as many endo-siRNA pathway proteins are known to localize to P granules, including the RNase III enzyme Dicer, the Dicer-related factor DRH-3, the endo-siRNA-binding Argonautes WAGO-1 and CSR-1, the piRNA-binding Argonaute PRG-1, and the RdRP EGO-1^20,23–25^. Other endo-siRNA pathway factors localize to other germline biomolecular condensates (WAGO-4 and ZNFX-1 in Z granules and RRF-1 in *Mutator* foci) that form ordered multi-condensate structures with P granules in adult *C. elegans* germ cells^22,26^. Together, the data hint that various *C. elegans* germ granules may act as organizational hubs that help connect endo-siRNA pathway proteins with the correct mRNAs to help ensure fidelity and accuracy of endo-siRNA-based gene regulation.

Here, we show that mutations that disrupt germ granule formation trigger the production of aberrant endo-siRNAs that inappropriately silence germline-expressed genes. Due to the heritable nature of endo-siRNAs, aberrations in gene expression are inherited across multiple generations, even after germ granules have been restored. We conclude that one function of germ granules is to organize and coordinate RNA-based epigenetic inheritance pathways and that the loss of this organizational function has consequences that can persist for many generations.

## Results

### *meg-3/4* animals exhibit defects in experimental RNAi

RNA interference (RNAi) can be triggered experimentally in *C. elegans* by feeding animals bacteria that express dsRNAs targeting specific *C. elegans* mRNA sequences (referred to hereinafter as experimental RNAi)^27^. Mutations in *deps-1* disrupt P granule formation in adult *C. elegans* germ cells and also lead to defects in experimental RNAi, suggesting that P granules may contribute in some way to small RNA-based gene regulation in germ cells^10^. To further investigate a potential link between P granules and germline small RNA pathways, we asked if other mutations that affect P granule formation also cause defects in experimental RNAi. MEG-3 and MEG-4 (together, MEG-3/4) are two intrinsically disordered proteins that are redundantly required for P granule assembly during early embryogenesis, but not during larval development or in the adult germline^8,9^. *tm4259* and *ax2026* are putative null alleles of *meg-3* and *meg-4,* respectively^8,28^. RNAi targeting either *pos-1* or *egg-4/5,* which are essential germline genes, causes sterility in *C. elegans* (Fig. 1A and Fig. S1A)^29^. *meg-3(tm4259) meg-4(ax2026)* animals remained fertile after either *pos-1* RNAi or *egg-5* RNAi, suggesting that MEG-3/4 are required for experimental RNAi in germ cells (Fig. 1A). *meg-3(tm4259) meg-4(ax2026)* animals also failed to silence a germline-expressed *gfp* reporter gene after exposure to *gfp* RNAi, confirming that MEG-3/4 contribute to RNAi-based gene silencing in germ cells (Fig. S1B). *meg-3(tm4259) meg-4(ax2026)* animals responded normally to RNAi targeting genes expressed in the soma, suggesting that the role of MEG-3/4 in promoting experimental RNAi may be restricted to the germline (Fig. S1C)^22^. *ax3055* and *ax3052* are independently isolated deletion alleles of *meg-3* and *meg-4*, respectively^9^. Similar to *meg-3(tm4259) meg-4(ax2026)* animals, *meg-3*(*ax3055*) *meg-4*(*ax3052*) animals responded normally to RNAi targeting somatically expressed genes, but failed to respond to RNAi targeting the germline-expressed *pos-1* and *egg-4/5* genes (Fig. 1A and Fig. S1A,C). The data show that MEG-3/4 contribute to dsRNA-based gene silencing in the germline and thereby support the idea that P granule formation is somehow important for dsRNA-based gene silencing.

**Figure 1.**
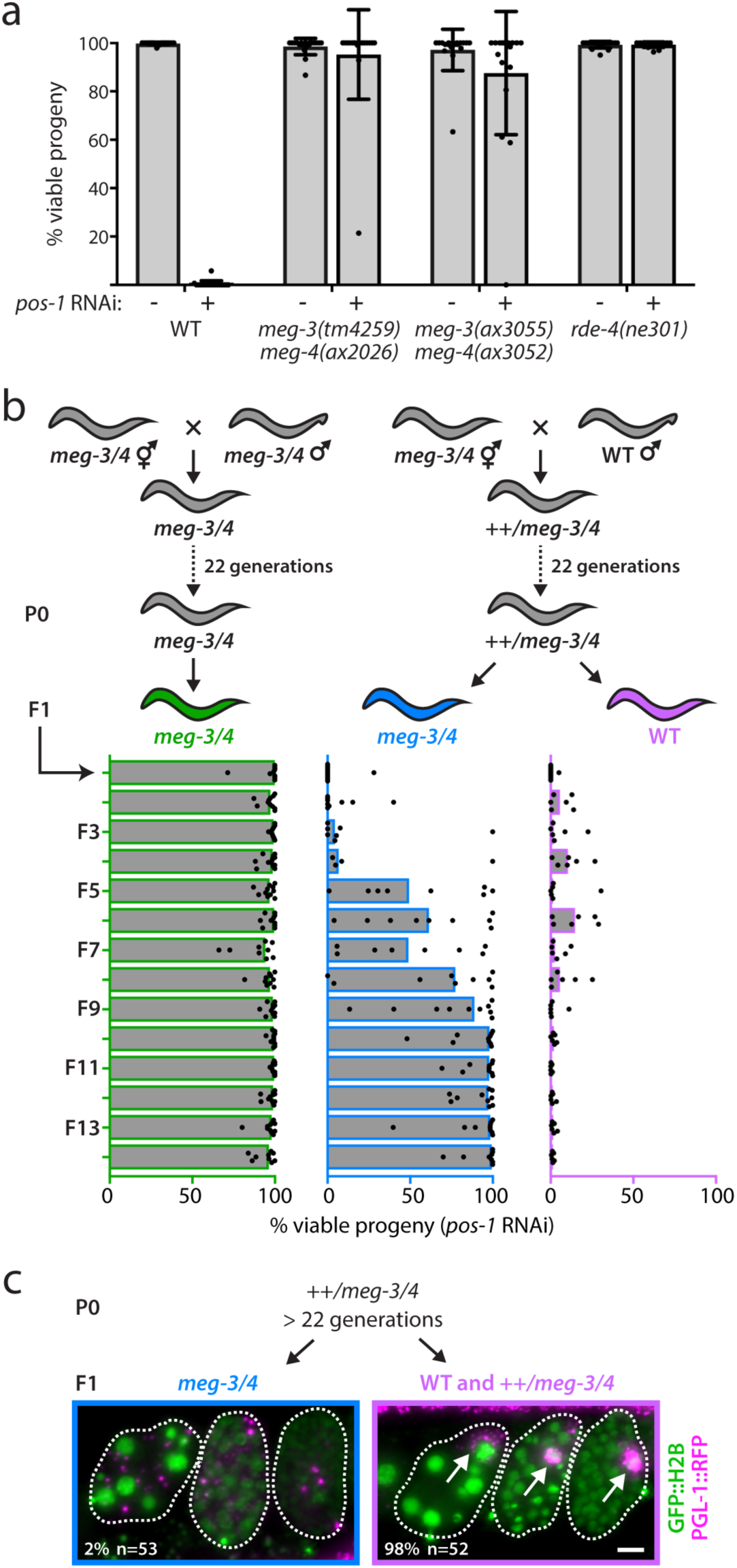
Transgenerational disconnect between *meg-3/4* genotype and phenotype. **(a)** L2 larvae of the indicated genotypes were fed either bacteria expressing dsRNAs derived from the *pos-1* gene, which is required for embryonic viability, or bacteria containing the control vector, L4440. When the animals became adults, they were allowed to lay broods, and % hatching embryos was scored. Black dots represent individual broods (n = 18). Error bars represent +/- standard deviations of the mean (gray bars). **(b)** Top panel: schematic of genetic crosses. We first marked *meg-3(tm4259) meg-4(ax2026)* with *dpy-3*(*e27)* (*dpy-3* is 0.1 cM from *meg-3* and 0.8 cM from *meg-4*). Then, *meg-3(tm4259) meg-4(ax2026) dpy-3(e27)* animals were crossed to wild-type males, and heterozygous progeny were identified by PCR-based genotyping of both *meg-3* and *meg-4*. As a mutant control cross, *meg-3(tm4259) meg-4(ax2026) dpy-3(e27)* animals were crossed to *meg-3(tm4259) meg-4(ax2026)* males. Lines that were either homozygous (control cross) or heterozygous for *meg-3/4* mutations were maintained for 22 generations (data presented later in the paper will clarify why this was necessary for this experiment). All lines remained heterozygous for *dpy-3(e27)*. The progeny of homozygous or heterozygous *meg-3/4* lines were isolated, and *meg-3* and *meg-4* were genotyped by PCR. These progeny were used to establish 6 lineages that were maintained for 14 generations under normal growth conditions. Bottom panel: at the indicated generations, animals from each lineage were exposed to *pos-1* RNAi and % hatching embryos was scored. Black dots represent individual lineages, colored bars represent the median value of % viable progeny. **(c)** A *meg-3(tm4259) meg-4(ax2026) dpy-3(e27)* chromosome was maintained in a heterozygous state for ≅45 generations in animals that were homozygous for two fluorescent protein markers: *pgl-1::rfp*^22^, which marks P granules (magenta), and *gfp::h2b*^31^, which marks chromatin (green). PGL-1::RFP and GFP::H2B were visualized in first-generation *meg-3/4* animals (indicated by Dpy phenotype) and their *meg-3/4(+)* and *++/meg-3/4* siblings (indicated by non-Dpy phenotype). Fluorescent micrographs of three embryos in the uterus of one adult are shown. Arrows indicate P granules. The percentage of F_1_ adults containing embryos with normal PGL-1::RFP expression and the number of adults scored are indicated. Scale bar, 10 microns. See also Figure S1.

### Transgenerational disconnect between *meg-3/4* genotype and phenotype

While conducting genetic crosses and RNAi experiments with *meg-3(tm4259) meg-4(ax2026)* animals (henceforth, *meg-3/4*), we noticed that the RNAi-defective (Rde) phenotype associated with *meg-3/4* was transgenerationally disconnected from the *meg-3/4* genotype. For instance, the *meg-3/4* offspring of heterozygous (++*/meg-3/4*) parents surprisingly responded to *pos-1* RNAi (Fig. 1B). Amazingly, descendants of these newly generated *meg-3/4* mutants did not become Rde until MEG-3/4 function had been absent for 5-9 generations (Fig. 1B). [Note: for reasons that will become apparent below, the ++*/meg-3/4* parents of the animals described above were maintained as heterozygotes for >20 generations prior to isolation of *meg-3/4* homozygous progeny in these experiments.] PGL-1 is a commonly used marker of P granules, and PGL-1 fails to concentrate into the germline blastomeres of MEG-3/4(-) embryos^8,9^. To ensure that MEG-3/4 function was lost in the *meg-3/4* animals described above, we monitored the subcellular localization of a PGL-1::RFP fluorescent marker protein in the various offspring of ++*/meg-3/4* parents. We found that PGL-1::RFP concentrated properly in wild-type and ++*/meg-3/4* embryos, but not in *meg-3/4* embryos (Fig. 1C). Importantly, the *meg-3/4* progeny in this experiment responded normally to RNAi, even though these animals did not possess detectable embryonic P granules (Fig. S1D). Thus, MEG-3/4 function is transgenerationally disconnected from Rde phenotypes. The fact that *meg-3/4* animals do not become Rde for many generations after the loss of MEG-3/4 indicates that MEG-3/4 do not play a direct role in dsRNA-based gene silencing. Rather, loss of MEG-3/4 triggers a process that indirectly impairs RNAi over many generations. Henceforth, we use the term “phenotypic lag” to refer to situations where a phenotype does not appear for many generations after genotype is established.

### Ancestral loss of P granules is associated with phenotypic hangovers

Outcrossing *meg-3/4* animals to wild type revealed a second type of transgenerational disconnect between *meg-3/4* genotype and phenotype. When *meg-3/4* animals were crossed to wild-type males, both the wild-type and *meg-3/4* F_2_ progeny of the cross were Rde (Fig. 2A,B). Amazingly, lineages established from the wild-type progeny of *meg-3/4* ancestors remained defective for *pos-1* RNAi for 9-11 generations before eventually reverting to a wild-type phenotype (Fig. 2B). Control crosses between wild type and wild type did not produce progeny exhibiting RNAi defects (Fig. 2A,B). Wild-type lineages derived from the *meg-3/4* outcross were also defective for RNAi targeting *egg-4/5* for >10 generations (Fig. S2A). Outcrossing a different set of *meg-3/4* alleles, *meg-3(ax3055)* and *meg-4(ax3052)*, also produced wild-type descendants that were RNAi-defective for multiple generations (Fig. S2B). The fact that wild-type descendents of *meg-3/4* animals retain an Rde phenotype for many generations indicates that mechanisms unrelated to *meg-3/4* genotype exist to propagate the Rde phenotype across generations. Henceforth, we use the term “phenotypic hangover” to refer to the transgenerational inheritance of a phenotype in the absence of the mutant genotype that originally triggered the phenotype. The remainder of this paper investigates the mechanism underlying *meg-3/4*-associated phenotypic hangovers.

**Figure 2.**
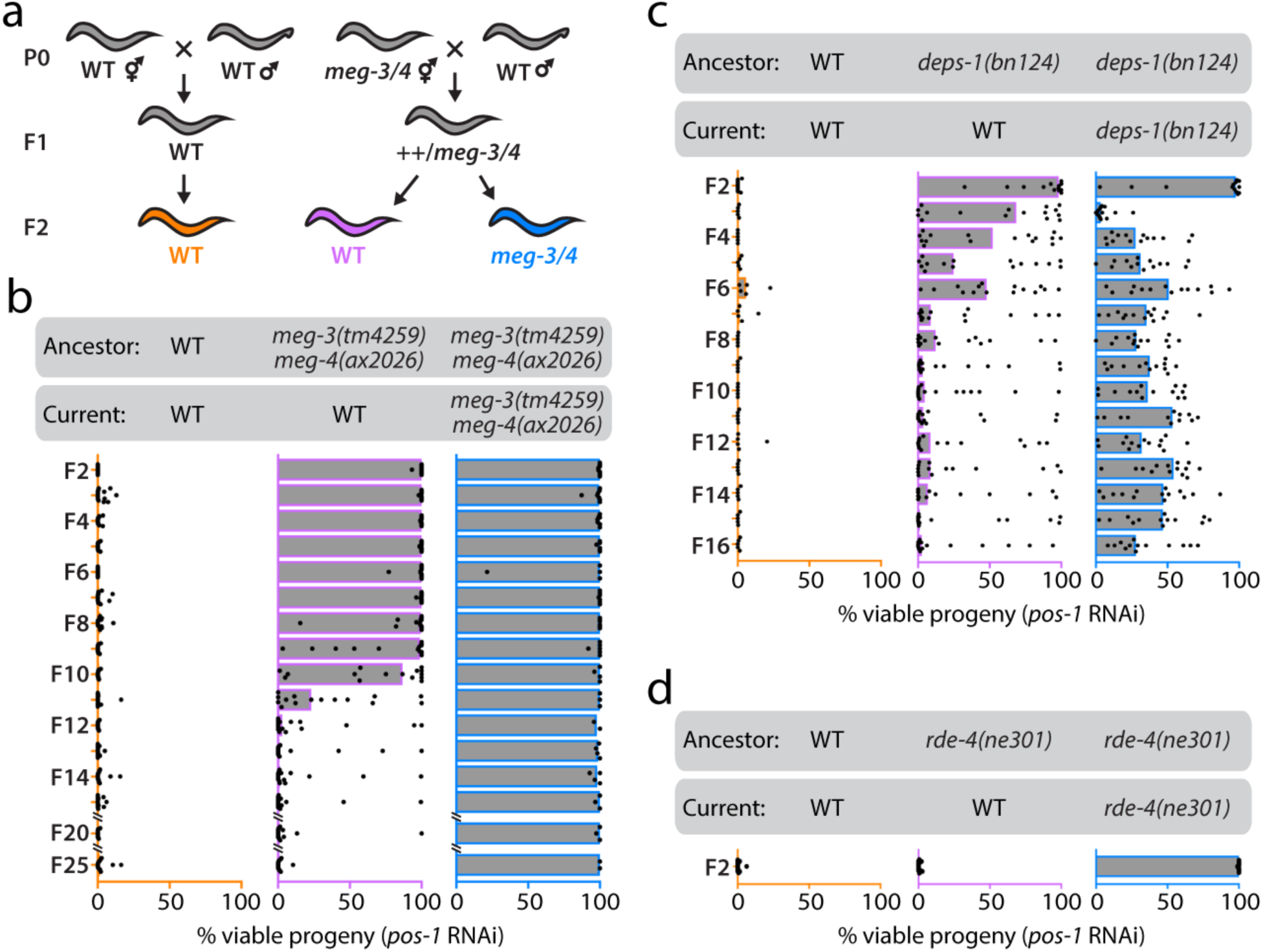
Ancestral loss of P granules is associated with phenotypic hangovers. **(a)** Schematic of genetic crosses used to generate animals scored for RNAi responsiveness in **(b)**. Note: for this experiment, *meg-3(tm4259) meg-4(ax2026)* animals had been maintained in a homozygous state for dozens of generations prior to outcross. **(b)** 15 lineages were established from F_2_ progeny and were maintained for 24 generations under normal growth conditions. At the indicated generations, animals from each lineage were exposed to *pos-1* RNAi and % hatching embryos was scored. Black dots represent individual lineages, colored bars represent the median value of % viable progeny. **(c)** *deps-1(bn124)* animals were crossed to wild-type males and descendants of the crosses were scored for *pos-1* RNAi sensitivity as described in **(b)**. **(d)** *rde-4(ne301)* animals were crossed to wild-type males, and wild-type and *rde-4* progeny were scored for *pos-1* RNAi sensitivity. See also Figure S2.

### Phenotypic hangovers are likely initiated by failure to assemble P granules

MEG-3/4 nucleate P granule assembly in germline blastomeres^8,9^. We wondered if the Rde hangovers associated with loss of MEG-3/4 might be related to the role of MEG-3/4 in P granule assembly. DEPS-1 is a *C. elegans* protein that 1) localizes to P granules, 2) is partially required for P granule assembly (primarily in adult germ cells), and 3) contributes to experimental RNAi in the germline^10^. To test our model that P granule disruption causes Rde hangovers, we asked whether mutations in *deps-1* could, like *meg-3/4*, trigger Rde hangovers. We crossed *deps-1(bn124)* animals to wild-type males, isolated homozygous wild-type or homozygous *deps-1* progeny, and measured RNAi responsiveness in lineages established from these animals. Both wild-type and mutant progeny of *deps-1(bn124)* ancestors were Rde, and lineages established from the wild-type progeny of *deps-1(bn124)* ancestors remained Rde for 2-5 generations after they had become genetically wild-type for *deps-1* (Fig. 2C). RDE-4 is a dsRNA-binding protein that is thought to play a direct role in dsRNA-mediated gene silencing by working with Dicer to process dsRNAs into siRNAs^30^. RDE-4 has no known role in P granule assembly. Unlike *meg-3/4* or *deps-1* outcrosses, *rde-4* outcrosses showed Mendelian inheritance of the Rde phenotype, indicating that (as expected) not all RNAi-related factors are associated with Rde hangovers (Fig. 2D). Altogether, our data suggest that Rde hangovers are likely triggered by an ancestral loss of P granules.

### The inheritance phase of Rde hangovers is not associated with obvious defects in germ granule morphology or localization

We wondered if Rde hangovers might be caused by inherited defects in P granule assembly. To test this idea, we first generated animals that were homozygous mutant for *meg-3/4* and that expressed PGL-1::RFP, a fluorescent marker of P granules^22^. We next outcrossed *pgl-1::rfp; meg-3/4* animals (after animals had been *meg-3/4* mutants for ≅10 generations) to *pgl-1::rfp; meg-3/4(+)* males and monitored PGL-1::RFP subcellular localization in embryos in the uteri of F_2_ progeny from this cross. Whereas embryonic P granules failed to concentrate properly in *meg-3/4* homozygous progeny, as expected, embryonic P granules formed properly in animals that had just become wild-type for *meg-3/4* (Fig. 3A). P granules also appeared normal in the adult germline in wild-type F_2_ progeny (Fig. S3A). We conclude that, although Rde hangovers are likely initiated by the loss of P granules, the maintenance of Rde hangovers is not associated with the inheritance of obvious defects in P granule morphology or localization.

**Figure 3.**
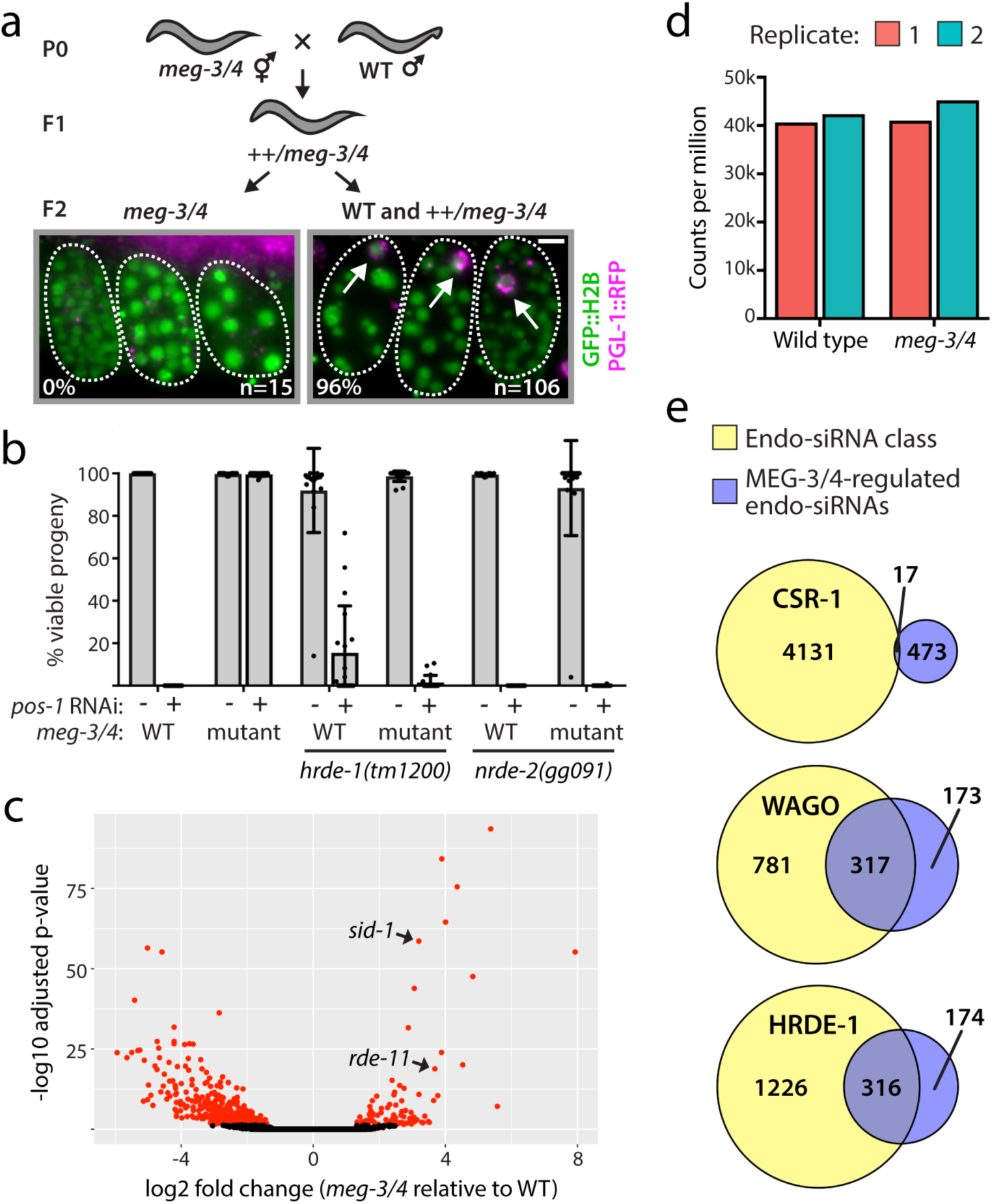
MEG-3/4 help organize endo-siRNA pathways. **(a)** *gfp::h2b; pgl-1::rfp; meg-3/4 dpy-3* animals that had been maintained in the homozygous state for dozens of generations were crossed to *gfp::h2b; pgl-1::rfp* males (WT). In the F_2_ generation, *meg-3/4(+)* and *++/meg-3/4* adults (indicated by non-Dpy phenotype) or *meg-3/4* homozygous adults (indicated by Dpy phenotype) were imaged. Fluorescent micrographs of three embryos in the uterus of one adult are shown. Arrows indicate P granules. The percentage of F_2_ adults containing embryos with normal PGL-1::RFP expression and the number of adults scored are indicated. Scale bar, 10 microns. **(b)** Loss of HRDE-1 or NRDE-2 suppresses the RNAi defect associated with *meg-3/4*. Individual animals of the indicated genotypes were scored for *pos-1* RNAi sensitivity as described in Figure 1A (n = 18). Error bars represent +/- standard deviations of the mean (gray bars). WT, wild type. **(c)** Volcano plot showing log2 fold change in the # of endo-siRNAs targeting each *C. elegans* gene in *meg-3(tm4259) meg-4(ax2026)* animals relative to wild type on the x-axis and the −log10 adjusted p-value on the y-axis. Dots shown in red indicate endo-siRNA pools that were significantly different between genotypes. *rde-11* and *sid-1* endo-siRNA pools are labeled for reasons that will become clear in Figure 5. **(d)** The number of endo-siRNAs targeting all MEG-3/4-regulated genes normalized to the total number of small RNAs sequenced from each sample are shown for each replicate of wild type and *meg-3/4*. k=1000. **(e)** Overlap between the list of MEG-3/4-regulated genes and published lists of genes targeted by CSR-1-bound endo-siRNAs^23^, genes in the WAGO class^20^, and genes targeted by HRDE-1-bound endo-siRNAs^18^. Numbers of genes overlapping and non-overlapping between lists are indicated. See also Figure S3.

### *hrde-1* suppresses the Rde phenotype associated with *meg-3/4* animals

Small RNAs are major vectors for transgenerational epigenetic inheritance (TEI) in plants and animals. Current models posit that, during endo-siRNA-directed TEI in *C. elegans*, a bolus of endo-siRNAs is deposited into the embryo via the egg or sperm. Parentally deposited endo-siRNAs then act as guide molecules to identify cognate mRNAs and recruit RdRP enzymes, which then amplify endo-siRNA populations. Finally, repetition of this process each generation allows gene regulatory information to pass across multiple generations. The germline-expressed Argonaute HRDE-1 is, for unknown reasons, required for endo-siRNA-directed TEI in *C. elegans*^18,19,31,32^. To ask if endo-siRNA-based TEI might somehow underlie Rde hangovers, we tested whether the Rde phenotype associated with *meg-3/4* animals depended on HRDE-1. Indeed, *tm1200*, a deletion allele of *hrde-1*, suppressed the RNAi defect, but not the P granule defect, associated with *meg-3/4* animals (Fig. 3B and Fig. S3B). HRDE-1 acts with Nuclear RNAi Defective-2 (NRDE-2) to drive endo-siRNA-based TEI^18,33^. NRDE-2 was also required for *meg-3/4* animals to exhibit an Rde phenotype (Fig. 3B). These data support the idea that endo-siRNA-directed TEI contributes in some way to the RNAi defect associated with *meg-3/4* animals.

### MEG-3/4 help organize endo-siRNA pathways

How might the endo-siRNA system contribute to RNAi defects and Rde hangovers? A number of endo-siRNA pathway factors localize to P granules (see Introduction). Thus, P granules may bring specific Argonautes, RdRPs, and mRNAs together in space and time in ways that help produce the correct numbers and types of endo-siRNAs each generation. This idea led us to the following model that might explain the mechanistic underpinnings of Rde hangovers. First, the disruption of P granules, via mutations like *meg-3/4* or *deps-1*, disorganizes endo-siRNA pathways, leading to aberrant production of endo-siRNAs targeting one or more genes required for RNAi (termed *RNAi gene-x* genes). Second, disorganized endo-siRNAs propagate across generations to transgenerationally silence *RNAi gene-x* (via HRDE-1), resulting in an Rde hangover. Note: for this model to work, *RNAi gene-x* would need to contribute to experimental RNAi, but not endo-siRNA-based TEI. To test our model, we sequenced small RNAs (≅15-30 nucleotides) using a 5’-phosphate-independent cloning method capable of sequencing *C. elegans* endo-siRNAs (see Method Details). We sequenced small RNAs from replicates of wild-type and *meg-3/4* animals, as well as genetically wild-type animals whose ancestors had been *meg-3/4* 3-25 generations prior to sequencing. Reads were mapped to the *C. elegans* genome, and the number of small RNAs mapping antisense to each *C. elegans* gene was quantified. We searched for genes that were differentially targeted by small RNAs in wild-type and *meg-3/4* animals (adjusted p-value < 0.05 and log2 fold change > 1 or < −1). The analysis identified 94 and 396 genes that were targeted by more or fewer small RNAs, respectively, in *meg-3/4* animals than in wild-type animals (termed MEG-3/4-regulated genes) (Fig. 3C and Table S1). Small RNAs targeting MEG-3/4-regulated genes were mostly 22 nucleotides in length, and the majority of 22-nucleotide RNAs initiated with guanosine (Fig. S3C). Given that such features are hallmarks of endo-siRNAs, we henceforth refer to these small RNAs as MEG-3/4-regulated endo-siRNAs^34^. Although the number of MEG-3/4-regulated endo-siRNAs went up or down in *meg-3/4* animals on a gene-by-gene basis, the total number of endo-siRNAs targeting all MEG-3/4-regulated genes remained similar in wild-type and *meg-3/4* animals (Fig. 3D). Thus, loss of MEG-3/4 alters the degree to which some genes are targeted by endo-siRNAs, but does not have a major effect on the overall number of endo-siRNAs produced. Previous studies have sub-categorized *C. elegans* endo-siRNAs into two major groups: endo-siRNAs associated with the Worm-specific Argonautes (WAGOs), and endo-siRNAs associated with the Argonaute CSR-1^20,23^. MEG-3/4-regulated endo-siRNAs were largely WAGO-class endo-siRNAs (p < 0.0001) (Fig. 3E). WAGO-class endo-siRNAs engage a number of WAGO-class Argonautes, including HRDE-1, to regulate gene expression^34^. MEG-3/4-regulated endo-siRNAs were enriched for HRDE-1-interacting endo-siRNAs (p < 0.0001) (Fig. 3E)^18^. Therefore, loss of MEG-3/4 and embryonic P granules is associated with aberrant levels of WAGO-class, HRDE-1-associated endo-siRNAs.

### Wild-type descendants of *meg-3/4* animals inherit aberrant endo-siRNA populations

Our model predicts that disruptions to endo-siRNA populations should persist in genetically wild-type descendants of *meg-3/4* animals for the duration of Rde hangovers (approximately ten generations). We therefore analyzed levels of MEG-3/4-regulated endo-siRNAs across generations in wild-type descendants of *meg-3/4* animals. Hierarchical clustering using the complete linkage method revealed three discernable patterns of inheritance (Fig. 4A). Most endo-siRNAs that were abundant in *meg-3/4* animals relative to the wild-type control remained high in the initial generations of the Rde hangover and then diminished to wild-type control levels over the course of 25 generations (Class 1 in Fig. 4A). Endo-siRNAs that were reduced in *meg-3/4* animals relative to the wild-type control fell into two major categories: a) endo-siRNAs that remained low in the initial generations of the hangover and increased slowly, many of which never fully recovered to wild-type control levels by the F_25_ generation (Class 2 in Fig. 4A); and b) endo-siRNAs that remained low in the initial generations of the hangover but recovered relatively quickly (Class 3 in Fig. 4A). Hierarchical clustering using divisive analysis also identified discrete patterns of endo-siRNA inheritance that resemble the categories described above (Fig. S4A). Both clustering analyses showed that the majority of MEG-3/4-regulated endo-siRNA pools progressed from mutant-like levels to wild-type levels over the course of 25 generations. A small proportion of MEG-3/4-regulated endo-siRNAs deviated from this general pattern of inheritance. For example, 6 pools of heritable endo-siRNAs remained at mutant-like levels for 25 generations and showed little evidence of reversion to wild-type levels (Fig. S4B). *hrde-1* affected the levels of 4 of these 6 endo-siRNA pools, suggesting that this remarkably stable inheritance was epigenetic (and not genetic) in nature (see Discussion). Interestingly, many aberrant endo-siRNA pools were unaffected by *hrde-1* in a *meg-3/4* mutant background, yet were still inherited in wild-type descendants of *meg-3/4* animals (Fig. S4C,D). Thus, HRDE-1-independent mechanisms of inheritance may exist in *C. elegans*.

**Figure 4.**
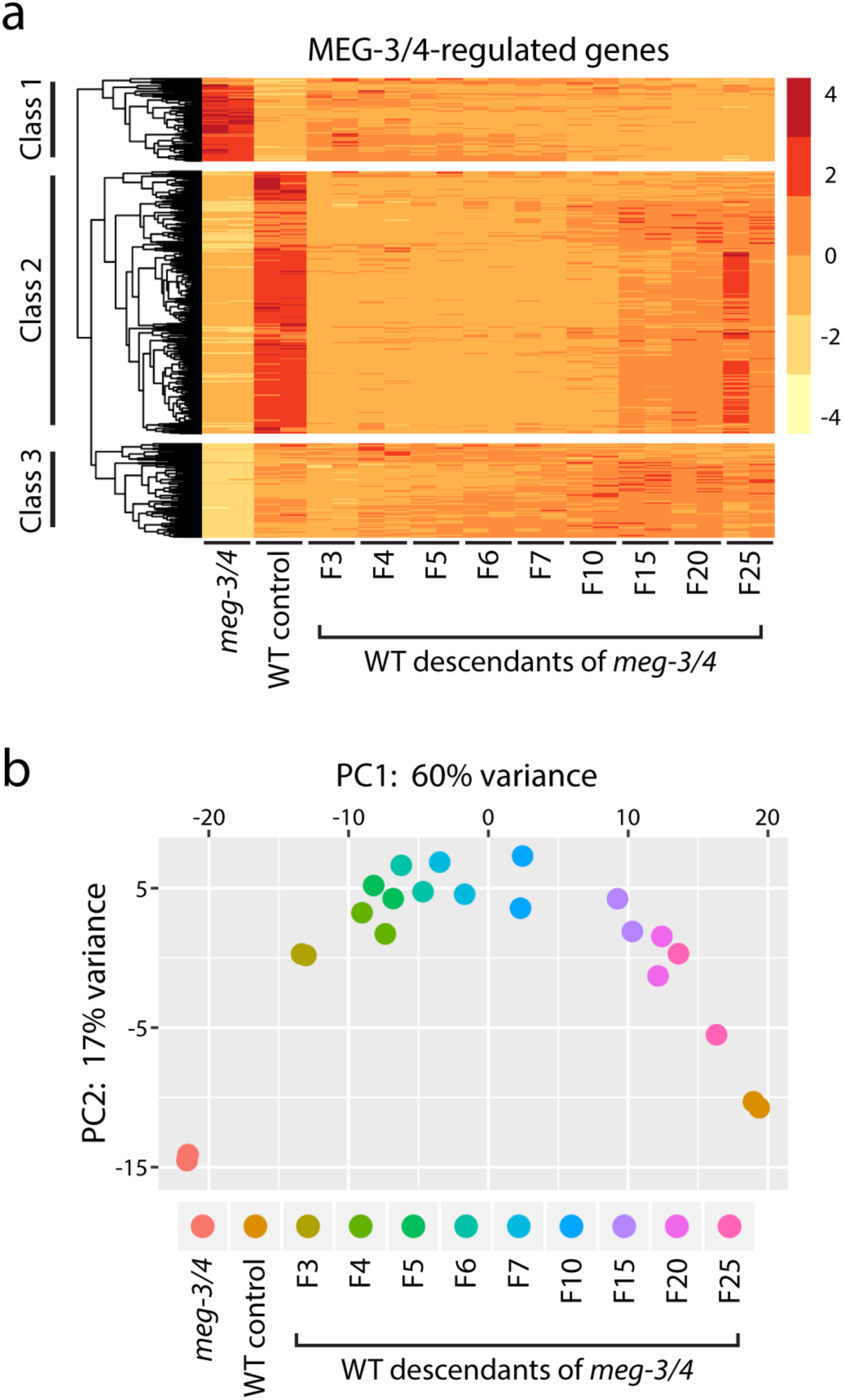
Wild-type descendants of *meg-3/4* inherit aberrant endo-siRNA populations. **(a)** Z scores of the levels of endo-siRNAs targeting each MEG-3/4-regulated gene. Biological replicates were plotted side-by-side for wild type (WT), *meg-3(tm4259) meg-4(ax2026) (meg-3/4),* and wild-type animals descending from *meg-3/4* mutant animals (P_0_) for the indicated number of generations (F_3_-F_25_). Genes were sorted into groups via hierarchical clustering (complete linkage method). The three major clusters of endo-siRNAs are shown and are indicated as Class 1-3. **(b)** Biplot of the top two principal components (PC1 and PC2) determined by principal component analysis. Points represent individual small RNA sequencing libraries, which are color-coded by genotype and by generation. Two biological replicates were analyzed for each genotype and each generation. Each PC represents a source of variation, and the distribution of points along a given PC axis indicates the degree of correlation between libraries for that PC. Note: PC1 explains most of the variation in the sequencing data. Prior to PCA, read counts were subjected to a regularized log transformation using the rlog function in DESeq2^51^. See also Figure S4.

The analyses described above indicate that aberrant endo-siRNA populations are inherited over many generations in animals whose ancestors lacked P granules. To further understand this remarkable pattern of inheritance, we subjected our sequencing data to principal component analysis (PCA). Briefly, PCA identifies and ranks sources of variation (termed principal components, or PCs) between datasets and then shows how the datasets relate to one another with regard to each PC. In the case of our small RNA sequencing data, for example, samples with similar small RNA profiles should cluster together in a plot of the highest-ranking PCs. PCA of our small RNA sequencing data revealed that the largest source of variation, PC1, corresponds to genotype, with *meg-3/4* animals and wild type exhibiting the least similarity (Fig. 4B). Interestingly, wild-type descendants of *meg-3/4* animals dispersed along the PC1 axis by generation: earlier generations more closely resembled *meg-3/4* animals, whereas later generations more closely resembled wild type, providing further evidence that wild-type descendants of *meg-3/4* animals evolve over the course of many generations from a mutant-like state to a wild-type state (Fig. 4B). PCA also revealed that, in each generation, biological replicates cluster together (Fig. 4B). Therefore, even though the recovery of endo-siRNA pools may take many generations, this process is reproducible and, therefore, largely deterministic.

### Ancestral loss of P granules is associated with heritable silencing of RNAi genes

Our model also predicts that aberrant endo-siRNA populations should inappropriately silence genes needed for experimental RNAi. Indeed, two of the top 12 most significantly upregulated endo-siRNA pools mapped to genes with known roles in experimental RNAi in *C. elegans* (*sid-1* and *rde-11*) (Fig. 3C and Table S1). SID-1 is a putative transmembrane protein that is expressed in the soma and germline, where it is thought to act as a channel for transporting dsRNA into cells^35–37^. RDE-11 is a zinc finger protein that is thought to function in the amplification step of RNAi^38,39^. *sid-1* and *rde-11* endo-siRNAs were elevated 9-fold and 13-fold in *meg-3/4* mutant animals, respectively, and remained elevated in wild-type descendants of *meg-3/4* animals for approximately 10 generations (Fig. 5A,B and Fig. S5). Thus, the generational kinetics of Rde hangovers and aberrant *sid-1/rde-11* endo-siRNA inheritance are quite similar. To determine the effects of these endo-siRNAs on gene expression, we used quantitative RT-PCR to measure the levels of *sid-1* and *rde-11* mRNA. *sid-1* and *rde-11* mRNA levels were down 5-fold and 6-fold in *meg-3/4* animals, respectively, and remained low in wild-type descendants of *meg-3/4* animals for approximately 10 generations (Fig. 5B and Fig. S5B). Heritable downregulation of *sid-1*/*rde-11* expression therefore also correlates with Rde hangovers. The data hint that runaway silencing of *sid-1* or *rde-11* (or both) may be responsible for *meg-3/4*-associated Rde hangovers. Since the RNAi defect associated with *meg-3/4* depends on HRDE-1, we asked whether the misregulation of *sid-1/rde-11* in *meg-3/4* animals also depends on HRDE-1. Indeed, *hrde-1* suppressed both the increased levels of *sid-1/rde-11* endo-siRNAs and the decreased levels of *sid-1/rde-11* mRNAs observed in *meg-3/4* animals (Fig. 5A,B and Fig. S5). Previous studies have shown that HRDE-1 interacts with endo-siRNAs targeting *sid-1* and *rde-11,* suggesting that the role of HRDE-1 in *sid-1/rde-11* silencing is likely to be direct^18^. Thus, HRDE-1 promotes both *sid-1/rde-11* silencing and germline Rde in the absence of MEG-3/4, and likely propagates these defects in wild-type animals whose ancestors lacked MEG-3/4. We conclude that ancestral loss of P granules is associated with heritable defects in gene expression, and that defects specific to *sid-1* and/or *rde-11* may be the cause of Rde hangovers.

**Figure 5.**
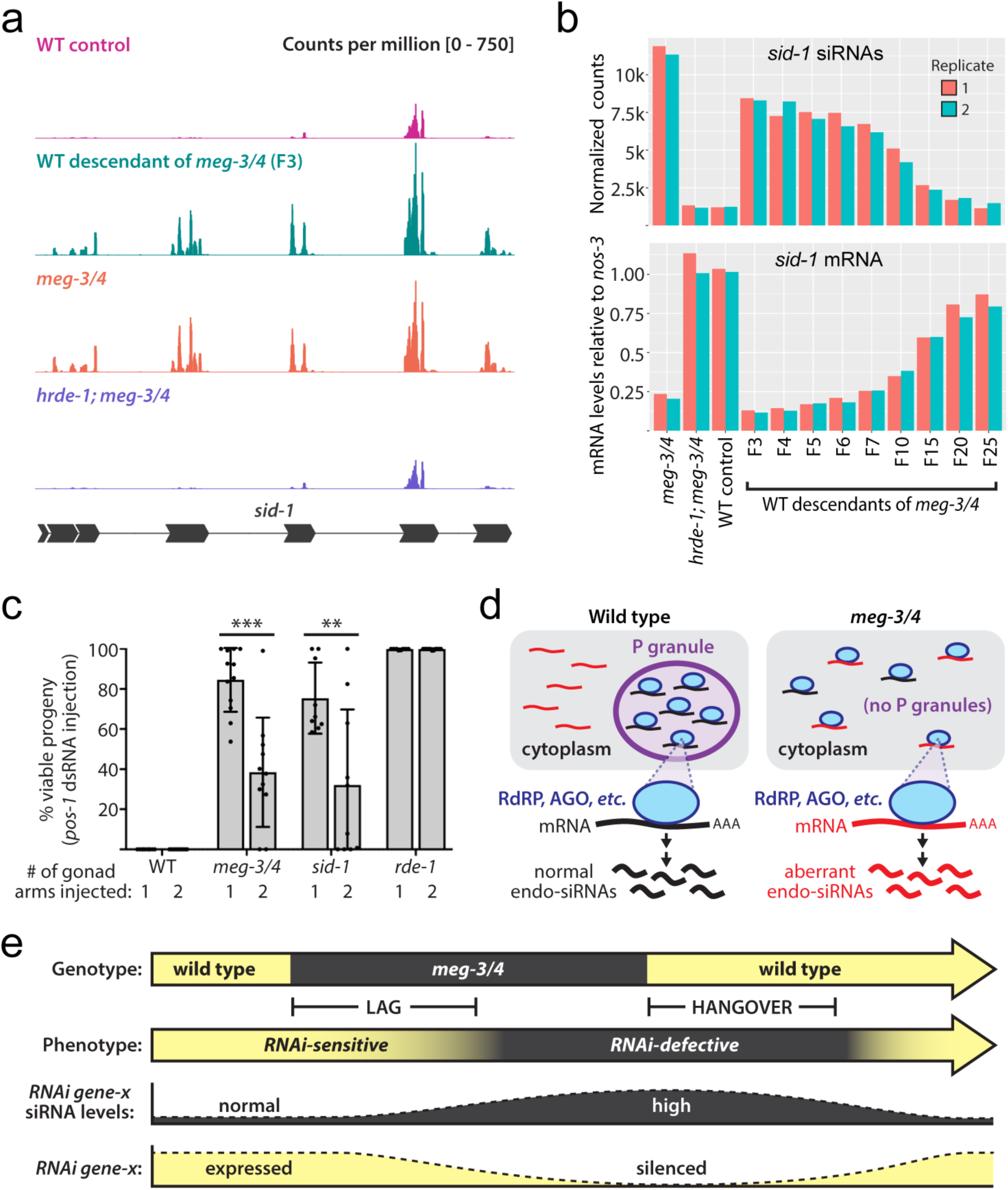
Ancestral loss of P granules is associated with heritable changes in *sid-1* expression. **(a)** endo-siRNAs sequenced from animals of the indicated genotypes (and generations after outcross) that map to the *sid-1* locus. A schematic of the *sid-1* locus is shown below. Counts were normalized to total number of reads. **(b)** Top panel: endo-siRNA reads mapping to the *sid-1* locus were quantified in replicates and across generations during Rde hangovers. Counts were normalized using the median of ratios method (DESeq2). Replicates are shown in red and blue. k=1000. Bottom panel: qRT-PCR was used to quantify *sid-1* mRNA in the samples shown in the top panel. *sid-1* mRNA values are shown relative to the mRNA values of *nos-3*, a germline-expressed gene. **(c)** Adults of the indicated genotype were injected with 200 ng/ul of *pos-1* dsRNA in either one or both arms of the germline (n ≥ 9 animals per condition). After 18-22 hours of recovery, animals were allowed to lay eggs and % hatching embryos was scored. Some of the variability observed in *meg-3/4* and *sid-1* animals may be due to the fact that responses of *sid-1* mutants to dsRNA injection change with time post-injection, and the timescale of this change varies from animal to animal^40^. Error bars represent +/- standard deviations of the mean (gray bars). **, p-value < 0.01; ***, p-value < 0.001 (Student’s t-test). WT, wild type. **(d)** Model for initiation of Rde hangovers: P granules coordinate and organize endogenous small interfering RNA (endo-siRNA) biogenesis by concentrating endo-siRNA pathway factors (blue ovals) such as RNA-dependent RNA polymerases (RdRPs) and Argonautes (AGOs) together with the appropriate mRNAs. In the absence of MEG-3/4 and therefore P granules, endo-siRNA pathway factors engage the “wrong” pool of mRNAs (long red lines) and thereby initiate the production of aberrant endo-siRNAs (short red lines). **(e)** Model to explain transgenerational disconnects between *meg-3/4* genotype and phenotype. Upon the loss of MEG-3/4, germ cells begin to produce slightly higher-than-normal levels of endo-siRNAs that target one or more genes required for experimental RNAi (*RNAi gene-x*). Due to the heritable nature of the endo-siRNA pathway, endo-siRNAs targeting *RNAi gene-x* accumulate slowly over generations in the absence of MEG-3/4 (lag). After 5-9 generations of aberrant endo-siRNA production/inheritance, these endo-siRNAs reach a level that causes silencing of *RNAi gene-x*; hence the defect in RNAi. After re-introduction of MEG-3/4, aberrant endo-siRNA pools continue to propagate across generations and continue to silence *RNAi gene-x* in genetically wild-type animals (hangover). Over the course of ten generations, genetic systems reassert their control over endo-siRNA biogenesis, returning endo-siRNA pools to normal. See also Figure S5.

*rde-11* and *sid-1* mutants respond idiosyncratically to dsRNAs injected directly into the germline^36,39,40^. For instance, *rde-11* mutants show a partial RNAi defect in response to low doses of injected dsRNA (0.5 ng/ul), but respond normally to high doses of injected dsRNA (20 ng/ul or 200 ng/ul)^39^. This differential sensitivity may be due to the presumed role of RDE-11 in amplifying gene silencing signals^38,39^. *sid-1* mutants, on the other hand, are sensitive to the number of gonad arms injected with dsRNA (the *C. elegans* germline is comprised of two distinct gonad arms)^36,40^. Because *C. elegans* RNAi is systemic (capable of spreading between tissues), dsRNA injection into a single gonad arm (or the soma) can trigger gene silencing in the germ cells of both gonad arms in wild-type animals (Fig. 5C)^41^. However, in *sid-1* mutants, which are defective for systemic RNAi, dsRNA injection only triggers gene silencing in the injected gonad and this silencing fails to spread to the other gonad arm^36,40^. We took advantage of these observations to ask if aberrant silencing of *rde-11* or *sid-1* might be the cause of Rde hangovers: We injected *meg-3/4* with a high dose of *pos-1* dsRNA (200 ng/ul) and asked if *meg-3/4* animals either 1) behaved like *rde-11* mutants (responded like wild-type to a high dose of dsRNAs), or 2) behaved like *sid-1* mutants (responded more robustly to injection of both gonad arms vs. a single arm). *meg-3/4* animals exhibited an Rde phenotype following injection of 200 ng/ul of *pos-1* dsRNA into either one or two gonad arms, suggesting that aberrant *rde-11* silencing is not solely responsible for the Rde phenotype of *meg-3/4* animals (Fig. 5C). Injecting both gonad arms of *meg-3/4* animals triggered more robust gene silencing than injection of a single gonad arm (Fig. 5C). *sid-1* mutant animals behaved similarly to *meg-3/4* animals in this assay (Fig. 5C). By contrast, *rde-1* mutant animals, which are fully defective for autonomous RNAi^42^, failed to respond to dsRNA injection regardless of the number of gonad arms injected (Fig. 5C). The data suggest that the RNAi defect associated with *meg-3/4* is largely systemic in nature. Thus, the data are consistent with a model in which the Rde phenotype associated with *meg-3/4* is caused by aberrant silencing of the *sid-1* gene. Note: for this model to be correct, aberrant silencing of the *sid-1* gene would need to be limited to germ cells, as *sid-1* mutant animals, but not *meg-3/4* mutant animals, exhibit RNAi defects in the soma (Fig. S1C)^36^.

## Discussion

Here, we show that mutations disrupting P granules trigger phenotypes that are transgenerationally disconnected from genotype. Disconnects correlate with alterations in endogenous small RNA levels and gene expression patterns, which persist across generations in the wild-type descendants of animals that lacked P granules. The data suggest that one function of germ granules is to organize and coordinate RNA-based modes of epigenetic inheritance. Upon loss of this germ granule-based organization, epigenetic defects are propagated on a generational timescale.

We find that MEG-3/4 regulate the degree to which particular genes are targeted by endo-siRNAs. How might MEG-3/4 (and therefore embryonic P granules) regulate endo-siRNA pathways? A subset of maternally deposited mRNAs and endo-siRNA pathway proteins (e.g. EGO-1, PRG-1, CSR-1, and WAGO-1) localize to P granules^13,20,23,24,43^. Thus, one function of P granules may be to spatially organize endo-siRNA biogenesis by ensuring that endo-siRNA pathway factors interact only with the proper mRNA targets (Fig. 5D). Such organization may be necessary to prevent the endo-siRNA system, which is inherently dangerous due to its feed-forward and heritable nature, from targeting functionally important germline genes for runaway heritable silencing. We propose that, in the absence of the organizing capabilities of P granules, one or more small RNA pathway factors fails to connect with its correct mRNA targets, resulting in over-targeting of some mRNAs and under-targeting of other mRNAs (Fig. 5D). The piRNA-binding Argonaute PRG-1 may be of particular relevance to this phenomenon, as PRG-1 shares a specific function in common with MEG-3/4: both PRG-1 and MEG-3/4 coordinate production of endo-siRNAs that bind HRDE-1^31,44,45^.

Because endo-siRNAs can be inherited, aberrant endo-siRNAs produced in the absence of P granules propagate for many generations, even after P granules have been restored. We find that wild-type descendants of *meg-3/4* ancestors can inherit aberrant endo-siRNA and mRNA patterns for over 25 generations. The fact that most endo-siRNA pools eventually return to wild-type levels (and do so reproducibly in biological replicates) indicates that, although recovery may take many generations, it is still largely a deterministic process. Thus, the information that ultimately dictates which mRNAs should and should not be channeled into the endo-siRNA pathway is likely hardwired in the genome. This hardwiring likely involves piRNAs, which are genomically encoded small RNAs with the ability to initiate endo-siRNA biogenesis in *C. elegans*^20,24,46,47^. Interestingly, we also find that some changes in endo-siRNA pools triggered by the loss of embryonic P granules were stably inherited for at least 25 generations. Epigenetic inheritance at these loci may represent situations in which endo-siRNA-based inheritance is so efficient that it has become largely divorced from genetic/piRNA control. Finally, we also identified a number of MEG-3/4-regulated endo-siRNA pools that were unaffected by *hrde-1*, hinting that HRDE-1-independent mechanisms for small RNA-based TEI may exist in *C. elegans*.

Our data argue against a direct role for embryonic P granules in experimental RNAi, as we have documented multiple examples of a) animals that lack embryonic P granules but respond normally to germline RNAi, and b) animals that possess P granules but fail to respond to germline RNAi. How, then, might *meg-3/4* mutations impair RNAi? A likely explanation is that the loss of P granules in *meg-3/4* triggers the production of endo-siRNAs that inappropriately and heritably silence one or more genes required for experimental RNAi in the germline (Fig. 5E). Multiple lines of evidence support this idea. First, *meg-3/4* animals produce abnormally high levels of endo-siRNAs that inappropriately silence *sid-1* and *rde-11*, two genes required for experimental RNAi. Second, aberrant levels of *sid-1*/*rde-11* endo-siRNAs and aberrant silencing of *sid-1*/*rde-11* mRNA are inherited for approximately 10 generations and, therefore, occur concomitantly with Rde hangovers. Third, aberrant *sid-1*/*rde-11* silencing and the *meg-3/4* RNAi defect both depend on the same factor: HRDE-1. Lastly, our dsRNA injection data point specifically to *sid-1* as the gene whose aberrant silencing triggers Rde hangovers, as *meg-3/4* mutant animals show a systemic defect in germline RNAi that mimics that of *sid-1* mutant animals. For the above reasons, we speculate that inappropriate and heritable silencing of the *sid-1* locus in germ cells is the cause of Rde hangovers. Proving this model will require additional work, which could include experimentally restoring *sid-1* expression to wild-type levels in animals undergoing Rde hangovers. A related model might explain the phenotypic lags we observe in *meg-3/4* animals. Specifically, loss of P granules (via introduction of *meg-3/4* mutations) might cause a small increase in *sid-1* endo-siRNAs that, initially, is not sufficient to effectively silence *sid-1*. In subsequent generations, however, *sid-1* endo-siRNAs would originate from two sources: 1) continued mild overproduction of endo-siRNAs (due to the absence of P granules), and 2) parental deposition of endo-siRNAs that were produced in previous generations. According to this model, over four to five generations, *sid-1* endo-siRNAs would accumulate to levels sufficient to silence the *sid-1* locus (Fig. 5E). Sequencing endo-siRNAs during a *meg-3/4* phenotypic lag would be a strong test of this idea.

In addition to *sid-1* and *rde-11*, hundreds of other genes are also mistargeted by aberrant endo-siRNAs in the wild-type descendants of *meg-3/4* animals (Fig. 4). It is possible that inappropriate silencing of these genes could trigger additional types of phenotypic lags or hangovers that have not yet been documented. Furthermore, mutations disrupting germ granule assembly or organization at other developmental time points might trigger a distinct set of genotype/phenotype disconnects, as the types of mRNAs that would be available to inappropriately enter the endo-siRNA pathway would likely change throughout development. Supporting this idea, some of the genes misregulated in *deps-1* animals (which exhibit defects in P granule assembly in both embryos and adult germ cells) were also identified by us in *meg-3/4* animals (which lack embryonic P granules), whereas other regulated genes were unique to each genotype (Table S2)^10^. Notably, *deps-1* mutations decrease the expression of *rde-4*, a gene required for RNAi; therefore, reduced levels of RDE-4 (and not SID-1) may underlie the germline RNAi defect of *deps-1* mutants^10^.

*C. elegans* possess a robust mode of TEI thought to transmit an RNA-based memory of germline gene expression programs from parent to progeny. We find that perturbing this mode of TEI (by disrupting P granule formation) leads to heritable defects in germline gene expression programs. Given that systems exist to transmit epigenetic information across generations, it stands to reason that genetic or environmental perturbations that alter the quality or quantity of this information would have heritable effects, even if the initiating perturbation were short-lived. Although heritable alterations to the epigenome could conceivably be adaptive, it is more likely that such changes would have negative impacts on organismal fitness, which could persist for generations. Such considerations may also apply to other modes of TEI and other animals, as mutations affecting chromatin or gene regulatory factors in both *C. elegans* and mammals have also been linked to phenotypic hangovers^48–50^. Given that many genes, pathways, and environmental signals are likely to impinge upon the germline epigenome, phenotypic lags and phenotypic hangovers, such as those documented here, may turn out to be a fairly common phenomenon. Exploring how much phenotypic variation is contingent upon ancestral genotype, assessing the generational perdurance of this type of variation in different animals, and asking if phenotypic hangovers ever contribute to disease inheritance in humans will be important questions for future studies to address.

## Acknowledgements

We thank the following people: members of the Kennedy lab for insights and suggestions, John Paul Ouyang and Geraldine Seydoux for sharing unpublished data, and Craig Hunter for suggesting the dsRNA microinjection experiment. Some strains were provided by the *Caenorhabditis* Genetics Center (CGC), which is funded by NIH Office of Research Infrastructure Programs (P40 OD010440). Some strains were provided by the Mitani laboratory through the National BioResource Project (Tokyo, Japan), which is part of the International *C. elegans* Gene Knockout Consortium. This work was supported by the National Institutes of Health, RO1GM088289 (S.K.). A.E.D is a Damon Runyon Fellow supported by the Damon Runyon Cancer Research Foundation (DRG-2304-17).

## Author Contributions

S.K and A.E.D conceived and performed experiments, wrote the manuscript, and secured funding.

## Materials and Methods

### *C. elegans* strains and growth conditions

*C. elegans* were grown at 20°C for all experiments. Unless otherwise indicated, animals were maintained on Nematode Growth Medium (NGM) plates seeded with *E. coli* OP50. Genetic crosses described in this study were between hermaphrodites and males. Phenotypic analyses were performed on hermaphrodites. A list of strains used in this study can be found in Table S3.

### Bacteria-mediated RNA interference

Animals were fed *E. coli* HT115 expressing dsRNAs targeting the indicated genes. *E. coli* HT115 containing the L4440 vector was used as a no-RNAi control in Figure 1A, Figure 3B, and Figure S1. RNAi clones, with the exception of *gfp* RNAi, were obtained from the Ahringer *C. elegans* RNAi library^52^. For tests using *pos-1* RNAi or *egg-5* RNAi, RNAi feeding began at the L2/L3 larval stages. *gfp* RNAi feeding began at the L3 larval stage, and *gfp* silencing was scored in the same generation at the adult stage. Somatic RNAi assays were performed by plating embryos onto bacteria expressing the indicated dsRNAs, and silencing was scored in the same generation at the adult stage. Details on scoring and sample size are described in figure legends.

### Phenotypic lag experiments

The *dpy-3* locus is approximately 0.1 cM from *meg-3* and 0.8 cM from *meg-4* (*meg-3* and *meg-4* are approximately 0.7 cM apart). We first marked a *meg-3(tm4259) meg-4(ax2026)* chromosome with *dpy-3(e27)*. Then, we crossed *meg-3(tm4259) meg-4(ax2026) dpy-3(e27)* animals to wild-type males and maintained a *meg-3(tm4259) meg-4(ax2026) dpy-3(e27)* chromosome in a heterozygous state for at least 22 generations. Multiple independent lines were established in this manner. Every generation, non-Dpy progeny were singled from non-Dpy parents that gave rise to both Dpy and non-Dpy progeny. Prior to performing lag experiments, lines were genotyped for *meg-3* and *meg-4* by PCR to check whether *meg-3(tm4259)* and *meg-4(ax2026)* were still linked to *dpy-3(e27)*. The progeny of animals that had been heterozygous for 22 generations were singled, tested for sensitivity to *pos-1* RNAi, and then genotyped for *meg-3* and *meg-4* by PCR (n = 72). In parallel, siblings of the progeny tested for RNAi were singled under normal growth conditions, genotyped for *meg-3* and *meg-4* by PCR, and wild-type and *meg-3/4* animals were used to establish lines (6 wild-type lines and 10 *meg-3/4* lines). Lines were maintained under normal growth conditions. Every subsequent generation, a small pool of animals (3-5) from each line was tested for sensitivity to *pos-1* RNAi. Lines were re-genotyped for *meg-3* and *meg-4* approximately every 5 generations. As a control, *meg-3(tm4259) meg-4(ax2026) dpy-3(e27)* animals were crossed to *meg-3(tm4259) meg-4(ax2026)* males, and the *dpy-3(e27)* allele was maintained in a heterozygous state for at least 22 generations, whereas *meg-3(tm4259)* and *meg-4(ax2026)* were maintained in a homozygous state. Fifteen *meg-3(tm4259) meg-4(ax2026) dpy-3(e27)* lines were then established and tested in parallel with the lines described in the first cross. To examine the P granule phenotype of newly generated *meg-3/4* mutants (Fig. 1C), similar crosses were performed as described above, except animals also contained *pgl-1(gg547[pgl-1::3xflag::tagrfp])* to mark P granules and *mjIs31[pie-1p::gfp::h2b]* to mark chromatin. For this experiment, *meg-3/4* genotype was inferred from the Dpy phenotype.

### Phenotypic hangover experiments

In general, hangover experiments began by crossing animals homozygous for the mutant allele(s) of interest [e.g. *meg-3(tm4259) meg-4(ax2026)*] to wild-type males (P_0_ generation). Multiple independent crosses were performed for each experiment. In the F_2_ generation, L2/L3 larvae were singled, tested for sensitivity to *pos-1* (or *egg-5*) RNAi, and then genotyped by PCR. Siblings of those animals were singled under normal growth conditions, genotyped by PCR, and were then used to establish either homozygous wild-type or homozygous mutant lines. Unless otherwise indicated, lines were maintained under normal growth conditions for the duration of the experiment. Each subsequent generation, a small pool of animals (3-5) from each line was tested for sensitivity to *pos-1* (or *egg-5*) RNAi. Lines were re-genotyped by PCR every 5-10 generations. As a control, wild-type animals descending from a cross between wild-type hermaphrodites and wild-type males were tested for sensitivity to *pos-1* (or *egg-5*) RNAi in parallel with the wild-type descendants of mutant animals. To examine the P granule phenotype in wild-type descendants of *meg-3(tm4259) meg-4(ax2026)* animals (Fig. 3A and Fig. S3A), similar crosses were performed as described above, except animals also contained *pgl-1(gg547[pgl-1::3xflag::tagrfp])* to mark P granules and *mjIs31[pie-1p::gfp::h2b]* to mark chromatin. In addition, the *meg-3(tm4259) meg-4(ax2026)* chromosome was marked with *dpy-3(e27)*, and *meg-3/4* genotype was inferred from the Dpy phenotype.

### Sample collection for small RNA-seq and qRT-PCR

The following crosses were performed in parallel in preparation for RNA isolation: 1) *meg-3(tm4259) meg-4(ax2026)* animals were crossed to wild-type males (P_0_ generation), and wild-type descendants of this cross were collected in generations F_3_ through F_7_, F_10_, F_15_, and F_25_; and 2) as a control, wild-type animals were crossed to wild-type males (P_0_ generation), and the descendants of this cross, which were all wild type, were collected in generation F_3_ (this sample is referred to as the wild-type control). For each type of cross, biological replicates were derived from different parents (P_0_s). To amass enough animals for RNA isolation, 30-50 wild-type lines were established in the F_2_ generation (for each biological replicate), and lines were pooled starting in the F_3_ generation. *meg-3(tm4259) meg-4(ax2026)* animals that had been homozygous mutant for dozens of generations were collected as a mutant control. Approximately 10% of *meg-3(tm4259) meg-4(ax2026)* animals do not develop a full germline, and 27% of *meg-3(tm4259) meg-4(ax2026)* animals are sterile (have empty uteri)^8^. To help control for the proportion of germ cells in each sample, adults with empty uteri were removed with a standard worm pick prior to sample collection. Adult worms were washed two times with M9 Buffer, resuspended and vortexed for 30 seconds in TRIzol, then flash-frozen in liquid nitrogen and stored at −80°C. Total RNA was isolated by TRIzol extraction.

### Small RNA library preparation and sequencing

RNAs ranging from approximately 15 to 30 nucleotides were gel-purified from total RNA (20 ug) on a 15% polyacrylamide/urea gel and then ligated to a 3’ adapter using T4 RNA ligase 2, truncated (New England BioLabs). To enable the cloning of 5’-triphosphorylated RNAs, samples were treated with Antarctic Phosphatase (New England BioLabs) followed by treatment with T4 polynucleotide kinase (New England BioLabs) as described previously^53^. Prior to ligation of the 5’ adapter, 3’-ligated small RNAs and any excess 3’ adapter were hybridized to the oligo that would eventually be used as a primer for reverse transcription. This step was taken to help minimize adapter-dimer formation^54^. To help avoid cross-contamination, the 5’ adapter was modified to contain the Illumina genomic sequencing primer annealing site followed by an additional 4 nucleotides at the 3’ end. Two different 5’ adapters (ending in either AGCG or CGUC) were mixed in a 1:1 ratio, and the mix was ligated to each sample using T4 RNA ligase I (New England BioLabs). Every sample was treated with the same mix of 5’ adapters. Libraries were amplified and multiplexed with a 6-nucleotide 3’ barcode, then pooled for next-generation sequencing on a NextSeq500 (Biopolymers Facility, HMS).

### Computational analysis of small RNA-seq

First, custom scripts were used to select reads starting with the last 4 nucleotides of the 5’ adapters (either AGCG or CGTC). Cutadapt 1.14 was used to trim the 3’ adapter (cutadapt -a CTGTAGGCACCATCAATAGATCGGAAGAGCAC -m 14 --discard-untrimmed) and the in-line portion of the 5’ adapter (cutadapt -u 4)^55^. Trimmed reads were then mapped to the *C. elegans* genome (WormBase release WS260) using Bowtie 1.2.2^56^. No mismatches were allowed. The number of reads mapping antisense to each gene was determined using featureCounts (featureCounts -s 2)^57^. Raw counts were then normalized by the median of ratios method using DESeq2 1.22.2 in R^51,58^. Differential analyses were performed using DESeq2 1.22.2^51^. Heatmaps (Fig. 4A and Fig. S4C) were clustered by row (genes) and scaled by row using pheatmap 1.0.12 in R^59^. The gene clusters shown in Figure S4A were generated using DEGreport 1.18.1 in R^60^. Principal component analysis of rlog-transformed counts (Fig. 4B) was performed using the rlog and plotPCA functions in DESeq2 1.22.2^51^. Custom scripts were used to extract the features of small RNAs that mapped antisense to MEG-3/4-regulated genes. Coverage plots (Fig. 5A and Fig. S5A) were generated as follows: first, bedGraph files normalized by counts per million were produced for the forward and reverse strands using deepTools 3.0.2 (bamCoverage -bs 5 --normalizeUsing CPM --samFlagExclude 16) and (bamCoverage -bs 5 --normalizeUsing CPM --samFlagInclude 16)^61^; then, bedGraph files were plotted in R using Sushi 1.20.0^62^.

### Quantitative RT-PCR

Using the total RNA prepared as described above, mRNA was reverse-transcribed into cDNA using the SuperScript III First-Strand Synthesis System (Invitrogen). Quantitative RT-PCR was performed with the iTaq Universal SYBR Green Supermix (Bio-Rad) and the primers listed in Table S4. Cycle threshold values were calibrated to a standard curve generated using a 4-point, 1:2 dilution series of wild-type control cDNA. PCR reactions were performed in technical triplicate for each biological replicate.

### Microscopy

Animals were immobilized in M9 Buffer containing 0.05% sodium azide and mounted on glass slides. Images were taken with a wide-field Zeiss Axio Observer.Z1 microscope equipped with an ORCA-Flash 4.0 CMOS camera (Hamamatsu) and the following Zeiss objectives: Plan-Apochromat 63×/1.4 Oil DIC M27, Plan-Apochromat 20×/0.8 M27, and EC Plan-Neofluar 10×/0.3 Ph1 M27. Embryos were imaged *in utero*. Images were acquired with ZEN software (Zeiss) and compiled in Fiji^63^.

### Synthesis and microinjection of *pos-1* dsRNA

*pos-1* dsRNA was synthesized *in vitro* using a MEGAscript T7 Transcription Kit (Invitrogen). The transcription template was PCR-amplified from the Ahringer *pos-1* RNAi clone and contained the *pos-1* insert sequence as well as the flanking T7 promoters. *pos-1* dsRNA was injected into one or both gonad arms of young adults at a concentration of 200 ng/ul. At least 9 animals were injected per condition. 18-22 hours post-injection, animals were singled and allowed to lay embryos for approximately 20 hours. Progeny and unhatched eggs were counted 24 hours later. A small fraction of *meg-3/4* animals did not lay eggs and were therefore excluded from the analysis. Mutant alleles were the following: *meg-3(tm4259) meg-4(ax2026)*, *sid-1(qt9)*, and *rde-1(ne219)*.

### Quantification and Statistical Analysis

Differential analyses for small RNA sequencing were performed with the Wald test using DESeq2 1.22.2, and genes with both an adjusted p-value < 0.05 and a log2 fold change >1 or < −1 were deemed significant^51^. Significance values reported in the description of Figure 3E were calculated with the one-sided Fisher’s exact test using the fisher.test function in R^58^. Significance values shown in Figure 5C were calculated with the Student’s t-test (two-tailed, unequal variances) using Excel. All error bars represent standard deviation. Sample sizes are indicated in the figure legends and Method Details.

