## Supplemental Information for "Germ Granules Coordinate RNA-based Epigenetic Inheritance Pathways"

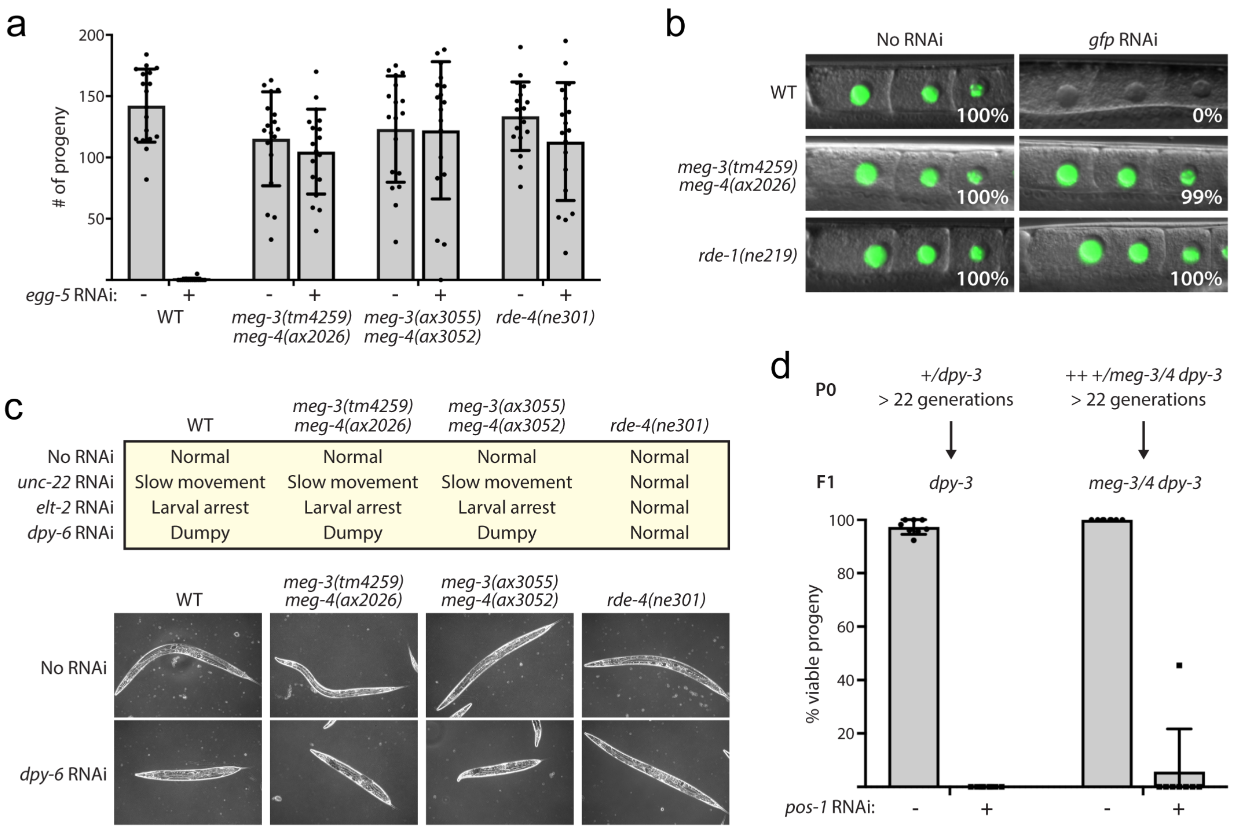


**Figure S1.** ***meg-3/4*** **animals are indirectly defective for germline RNAi, related to Figure 1. (a)** *meg-3/4* animals fail to respond to RNAi targeting *egg-4/5*. EGG-4/5 are two redundant, phosphatase-like proteins required for egg shell formation in fertilized *C. elegans* embryos[29]. Animals exposed to dsRNA targeting *egg-5* silences both *egg-4* and *egg-5* in adult *C. elegans*, causing animals to not produce any viable progeny[29]. L2 larvae of the indicated genotypes were fed either bacteria expressing dsRNAs derived from the *egg-5* gene or bacteria containing the control vector, L4440. When animals became adults, they were allowed to lay eggs for approximately 22 hours, and the number of progeny was scored 24 hours later. Black dots represent individual broods (n = 18). Error bars represent +/- standard deviations of the mean (gray bars). **(b)** *meg-3/4* animals containing a germline-expressed *gfp::h2b* transgene fail to silence *gfp* after exposure to *gfp* RNAi. Starting in the L3 larval stage, animals were fed bacteria expressing dsRNAs targeting *gfp*. After 24 hours of feeding on *gfp* RNAi (or a no-RNAi control), adults were imaged for *gfp* expression. Micrographs show GFP fluorescence in oocytes. The percentage of fluorescent animals is indicated (n = 100). **(c)** *meg-3/4* animals respond to somatic RNAi. Top panel: summary of responses to RNAi targeting either *unc-22*, *elt-2*, or *dpy-6*. Eggs were plated onto bacteria expressing dsRNAs targeting the indicated genes. After 3 days, adults were scored for the indicated phenotypes. Scoring was performed blind and on a plate-by-plate basis (12 plates per condition with approximately 100 animals per plate). For each condition, the result was the same for all 12 plates (e.g. 12/12 plates of wild-type animals scored "Dumpy" in response to *dpy-6* RNAi). Bottom panel: representative images of the indicated genotypes after feeding on *dpy-6* RNAi (or a no-RNAi control). **(d)** Siblings of the animals imaged in Figure 1C were tested for Rde, and *meg-3/4* animals were responsive to *pos-1* RNAi. A *meg-3(tm4259) meg-4(ax2026) dpy-3(e27)* chromosome was maintained in the heterozygous state as described in Figure 1C of the main text. Animals were homozygous for *gfp::h2b* and *pgl-1::rfp*. First-generation *meg-3/4* progeny (indicated by Dpy phenotype) were scored for *pos-1* RNAi sensitivity as described in Figure 1A. As a control, the *dpy-3(e27)* allele was maintained in the heterozygous state in *gfp::h2b; pgl-1::rfp; meg-3/4(+)* animals for an equal number of generations, and Dpy *meg-3/4(+)* animals were scored for *pos-1* RNAi sensitivity. Black dots represent individual broods (n = 8). Error bars represent +/- standard deviations of the mean (gray bars).

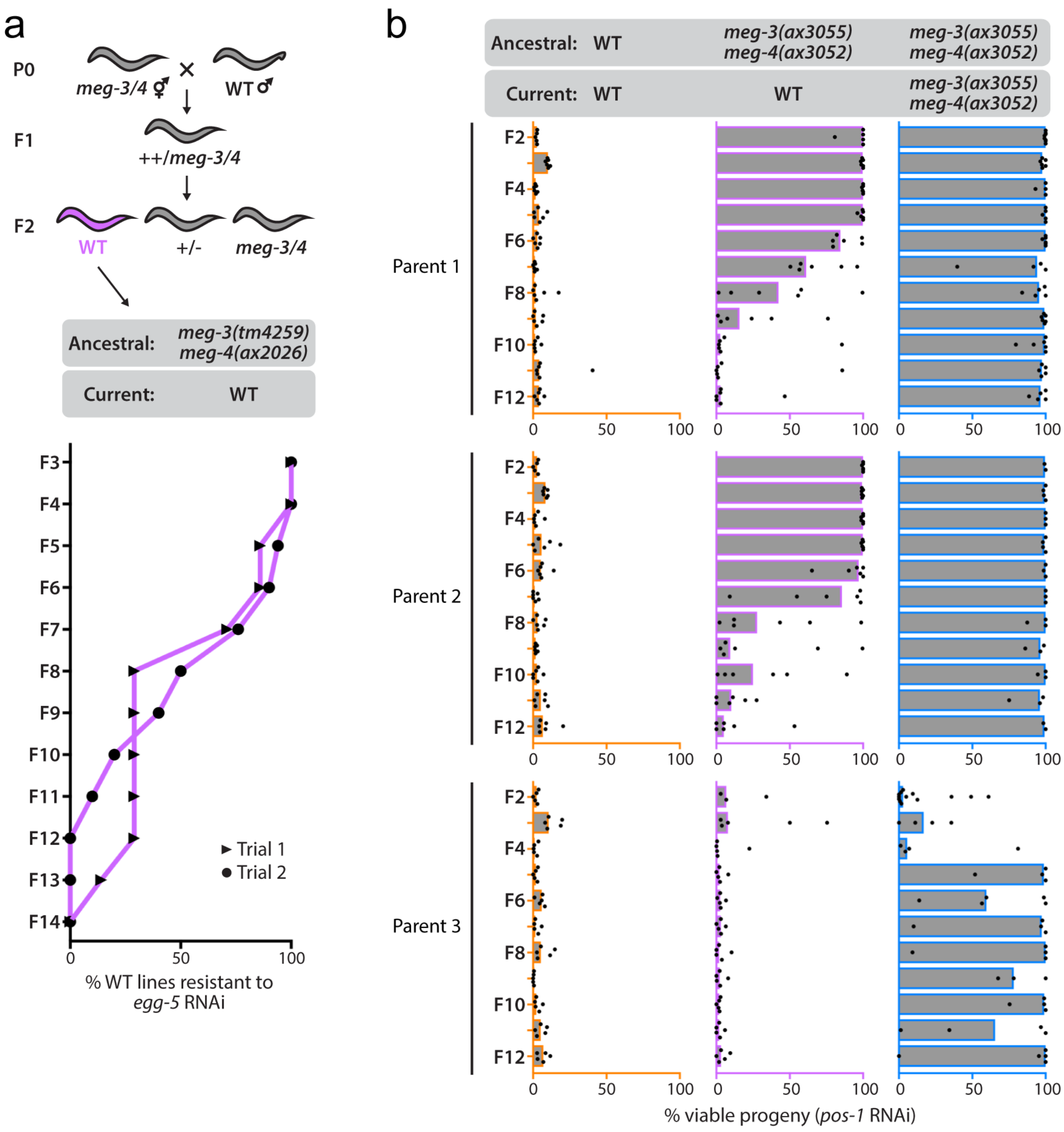


**Figure S2. Additional phenotypic hangovers triggered by P granule loss, related to Figure 2. (a)** Wild-type descendants of *meg-3/4* animals inherit defects in RNAi against *egg-4/5*. Top panel: *meg-3/4* animals were crossed to wild-type males as described in Figure 2A of the main text. Wild-type progeny of *meg-3/4* animals were isolated, and lineages were established from these animals. Bottom panel: each generation, animals were scored for resistance to *egg-5* RNAi. As described in Figure S1A, dsRNAs derived from the *egg-5* gene target both *egg-4* and *egg-5*[29]. In trial 1, wild-type lines were maintained under normal growth conditions. For each line, 5 L2/L3 larvae were pooled and tested for RNAi responsiveness every generation, which is a similar experimental setup as that used to generate the data in Fig. 2B of the main text. Lines that produced 10 or more viable progeny in response to *egg-5* RNAi were classified as resistant to *egg-4/5* RNAi. In trial 2, animals were maintained under *egg-5* RNAi conditions throughout the duration of the experiment. Wild-type lines that survived under these conditions were classified as resistant to *egg-4/5* RNAi. Note: in addition to showing an Rde hangover for *egg-4/5* RNAi, these data also show that putting animals under strong selection for inheriting Rde (animals are sterile if they do not inherit Rde) does not affect the generational perdurance of an Rde hangover. **(b)** An independently isolated set of *meg-3/4* alleles triggers Rde hangovers. Rde hangovers for *meg-3(ax3055) meg-4(ax3052)* animals were assessed as described for *meg-3*(tm4259) *meg-4*(ax2026) animals in Figure 2A,B of the main text. Black dots represent individual lineages, colored bars represent the median value of % viable progeny. Note that one of the three *meg-3(ax3055) meg-4(ax3052)* parents (Parent 3) produced offspring that did not show an Rde phenotype or an Rde hangover. This may be related to the observation that a small percentage (approximately 5-10%) of *meg-3(tm4259) meg-4(ax2026)* individuals do not exhibit an Rde phenotype (see Fig. 1A and Fig. S1A). Assuming that aberrant silencing of *sid-1* underlies the RNAi defect of *meg-3/4* mutants, the variability of Rde levels (and Rde hangovers) in *meg-3(ax3055) meg-4(ax3052)* animals may be due to variable levels of *sid-1* endo-siRNAs amongst animals lacking P granules. Consistent with this idea, the *meg-3/4* descendants of Parent 3 were sensitive to *pos-1* RNAi in the first three generations. This pattern of inheritance was reminiscent of the phenotypic lag described in Figure 1B of the main text and hints that all the progeny of Parent 3, regardless of their *meg-3/4* genotype, inherited levels of *sid-1* endo-siRNAs that were insufficient to cause an Rde phenotype.

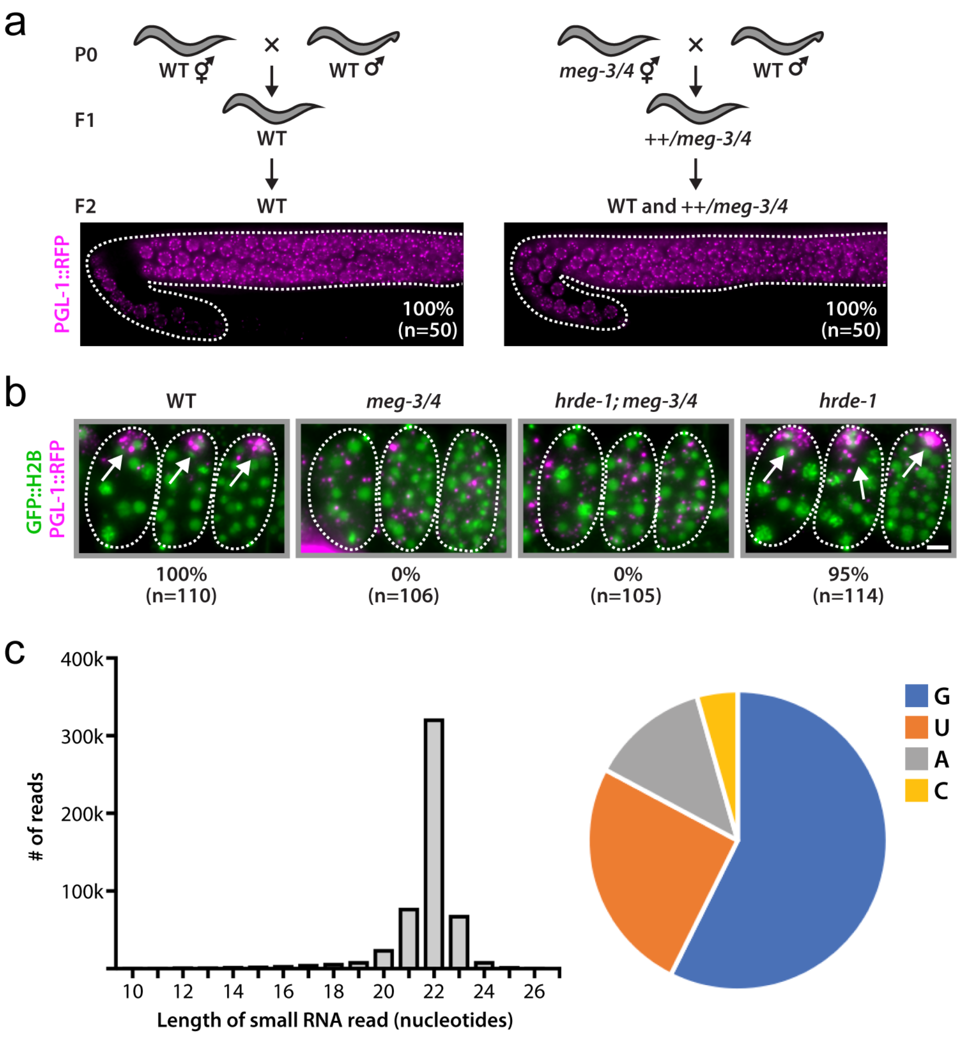


**Figure S3. Loss of P granules alters endo-siRNA pathways, related to Figure 3. (a)** P granules continue to form in the adult germline after a *meg-3/4* outcross. Either *gfp::h2b; pgl-1::rfp; dpy-3* animals (WT) or *gfp::h2b; pgl-1::rfp; meg-3/4 dpy-3* animals (*meg-3/4*) were crossed to *gfp::h2b; pgl-1::rfp* males (WT). In the $F_2$ generation, *meg-3/4(+)* and ++*/meg-3/4* adults (indicated by non-Dpy phenotype) were imaged for *pgl-1::rfp* expression. The percentage of $F_2$ adults containing germlines with normal PGL-1::RFP expression and the number of adults scored are indicated. Dotted line outlines the adult germline. **(b)** *hrde-1* does not suppress the P granule defect associated with *meg-3/4*.

Fluorescent micrographs show three embryos *in utero* for the indicated genotypes. PGL-1::RFP (magenta) marks P granules (arrows) and GFP::H2B (green) marks chromatin. The percentage of animals containing embryos with normal PGL-1::RFP expression and the number of animals scored are indicated. Scale bar, 10 microns. **(c)** MEG-3/4-regulated small RNAs are predominantly 22G endo-siRNAs. Left panel: length distribution of small RNAs that mapped antisense to MEG-3/4-regulated genes. k=1000. Right panel: pie chart of 22-nucleotide small RNAs that mapped antisense to MEG-3/4-regulated genes, grouped by the identity of the first nucleotide.

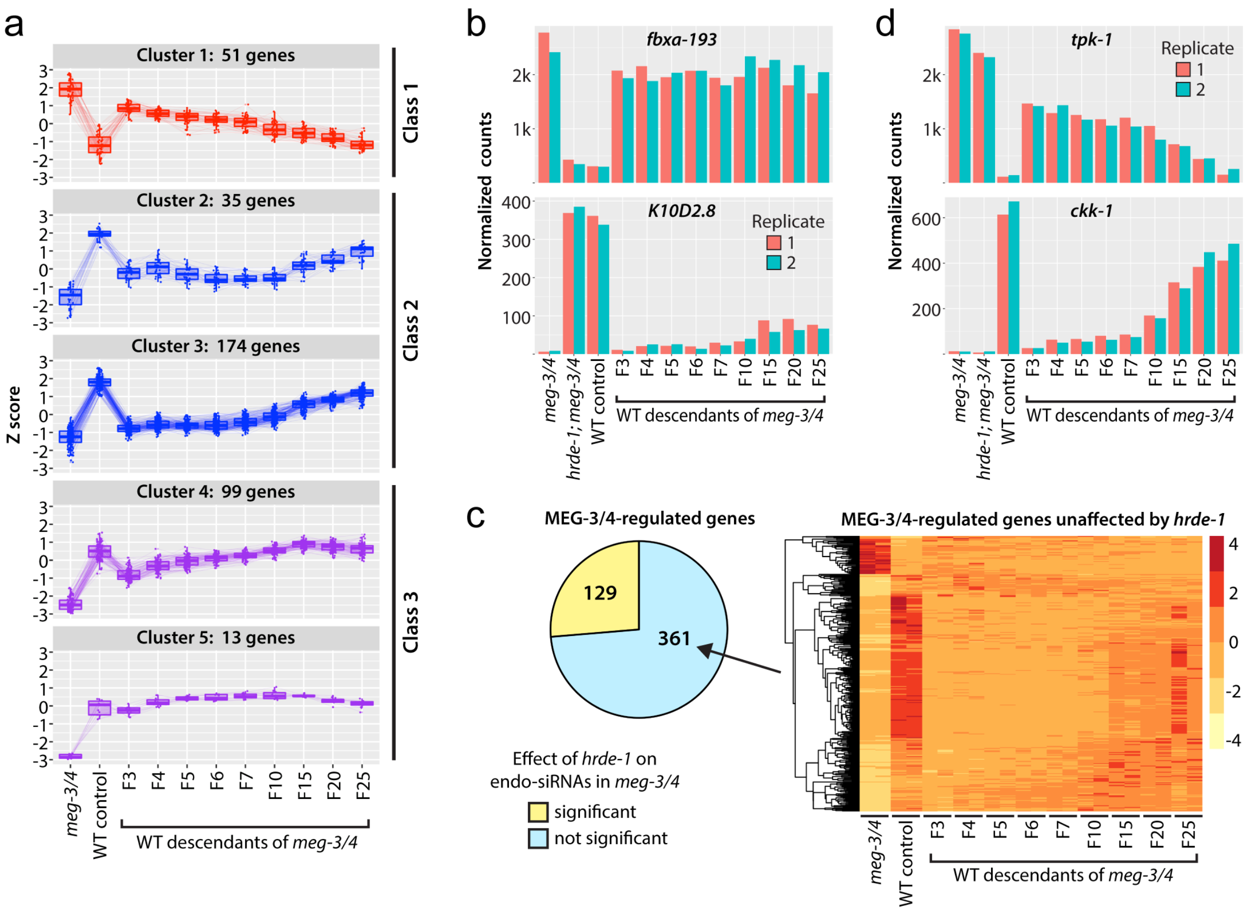


**Figure S4. Inheritance patterns of aberrant endo-siRNA populations in wild-type descendants of *meg-3/4*, related to Figure 4. (a)** MEG-3/4-regulated genes grouped by endo-siRNA inheritance patterns. Genes were sorted into groups via hierarchical clustering (DEGreport, divisive analysis method). Groups containing 10 or more genes are shown. The clustering method revealed five major clusters, which exhibited three general patterns of inheritance that were similar to Figure 4A of the main text. Class 1: endo-siRNA pools that were upregulated in *meg-3/4* animals and gradually decreased in wild-type descendants over the course of 25 generations. Class 2: two clusters of endo-siRNA pools that were downregulated in *meg-3/4* animals and recovered relatively slowly over the course of 25 generations. Class 3: two clusters of endo-siRNA pools that were downregulated in *meg-3/4* animals and recovered relatively quickly. **(b)** Two examples of MEG-3/4-regulated endo-siRNA pools that did not recover to wild-type levels by the $F_{25}$ generation of the hangover. Endo-siRNA reads mapping to the *fbxa-193* and *K10D2.8* loci were quantified in replicates and across generations during Rde hangovers. Counts were normalized using the median of ratios method (DESeq2). Replicates are shown in red and blue. k=1000. Six genes in total were identified as targets of endo-siRNAs that did not recover to near-wild-type levels. To identify such genes, we first searched for genes that were differentially targeted by small RNAs in wild-type $F_{25}$ descendants of *meg-3/4* animals (the last generation of the hangover that was tested) and wild-type control animals (adjusted p-value < 0.05 and log2 fold change > 1 or < -1). This analysis identified 11 gene targets, 6 of which overlapped with the list of MEG-3/4-regulated gene targets. **(c)** Heritable pools of aberrant endo-siRNAs unaffected by *hrde-1*. Left panel: proportion of MEG-3/4-regulated endo-siRNA pools that were affected by *hrde-1*. We asked which MEG-3/4-regulated endo-siRNA pools were dependent on HRDE-1 by first searching for genes that were differentially targeted by small RNAs in *meg-3/4* and *hrde-1; meg-3/4* animals (adjusted p-value < 0.05 and log2 fold change > 1 or < -1). We then used the results of this analysis to ask, for each MEG-3/4-regulated gene, whether its associated endo-siRNA pools were significantly affected by *hrde-1* in a *meg-3/4* background. Right panel: Z scores of MEG-3/4-regulated endo-siRNA pools that were unaffected by *hrde-1*. Biological replicates were plotted side-by-side for wild type (WT), *meg-3(tm4259) meg-4(ax2026) (meg-3/4),* and wild-type animals descending from *meg-3/4* mutant animals ($P_0$) for the indicated number of generations ($F_3$-$F_{25}$). Genes were sorted into groups via hierarchical clustering (complete linkage method). **(d)** Two examples of aberrant MEG-3/4-regulated endo-siRNA pools that were inherited over the course of the Rde hangover but did not show dependence on *hrde-1* in a *meg-3/4* background. Endo-siRNA reads mapping to the *tpk-1* and *ckk-1* loci were quantified in replicates and across generations during Rde hangovers. Counts were normalized using the median of ratios method (DESeq2). Replicates are shown in red and blue. k=1000.

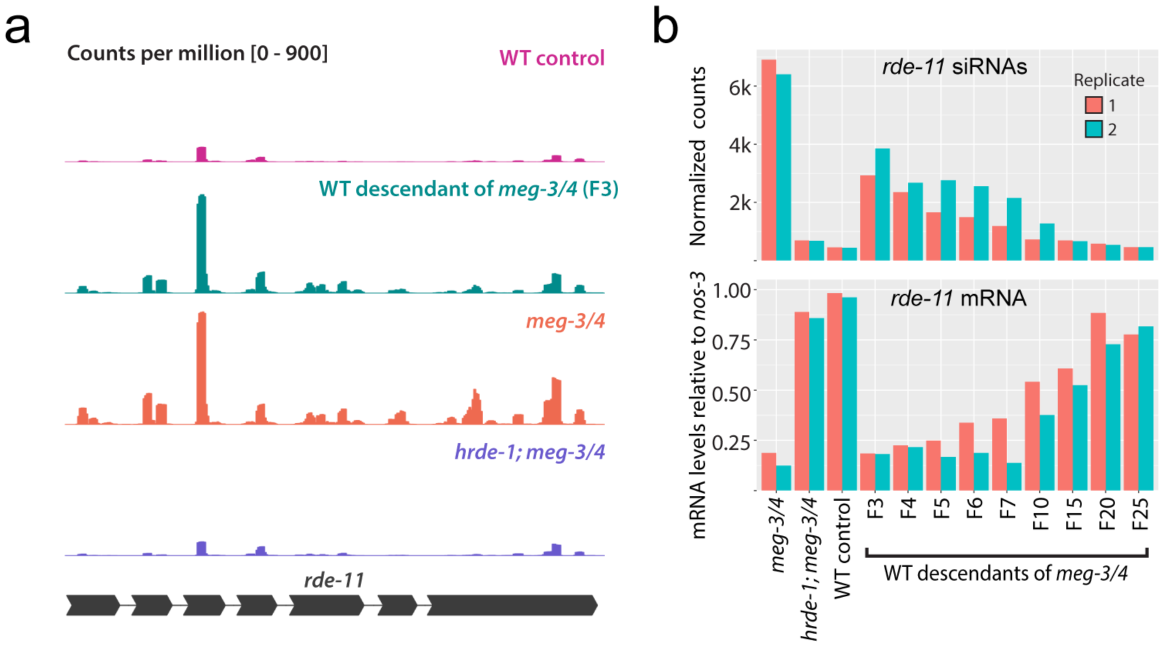


**Figure S5. Transgenerational silencing of *rde-11* correlates with Rde hangovers, related to Figure 5. (a)** Endo-siRNAs sequenced from animals of the indicated genotypes (and generations after outcross) mapping to the *rde-11* locus. A schematic of the *rde-11* locus is shown below. Counts were normalized to the total number of reads. **(b)** Top panel: endo-siRNA reads mapping to the *rde-11* locus were quantified in replicates and across generations during Rde hangovers. Counts were normalized using the median of ratios method (DESeq2). Replicates are shown in red and blue. k=1000. Bottom panel: qRT-PCR was used to quantify *rde-11* mRNA in the samples shown in the top panel. *rde-11* mRNA values are shown relative to the mRNA values of *nos-3*, a germline-expressed gene.

**Table S1. List of MEG-3/4-regulated endo-siRNA targets.**

**Table S2. Loss of DEPS-1 and MEG-3/4 affect a partially overlapping set of genes.**

**Table S3. List of strains used in this study.**

**Table S4. List of oligos used in this study.**
